# A refined pH-dependent coarse-grained model for peptide structure prediction in aqueous solution

**DOI:** 10.1101/2022.12.20.521219

**Authors:** Pierre Tuffery, Philippe Derreumaux

## Abstract

Peptides carry out diverse biological functions and the knowledge of the conformational ensemble of polypeptides in various experimental conditions is important for biological applications. All fast dedicated softwares perform well in aqueous solution at neutral pH. In this study, we go one step beyond by combining the Debye-Hückel formalism for charged-charged amino acid interactions and a coarse-grained potential of the amino acids to treat pH and salt variations. Using the PEP-FOLD framework, we show that our approach performs as well as the machine-leaning AlphaFold2 and TrRosetta methods for 15 well-structured sequences, but shows significant improvement in structure prediction of six poly-charged amino acids and two sequences that have no homologous in the Protein Data Bank, expanding the range of possibilities for the understanding of peptide biological roles and the design of candidate therapeutic peptides.

## 2 Introduction

Peptides are short polymers of few tens of amino acids that carry out diverse biological functions, acting as signaling entities in all domains of life, targeting receptors or interfering with molecular interactions. Hormones and their bacterial mimetics [1], neuropeptides and their roles in neurodegenerative diseases [2], antimicrobial peptides contribution to host defence [3], immunomodulatory peptides in the perspective of vaccine design [4], are some current directions motivating their study at a fundamental level. However, due to their specific features, peptides have also gained interest as therapeutical agents [5], particularly to target protein-protein interactions [6]. They are also considered as having potential to develop new functional biomimetic materials [7]. Short peptides have limitations though, such as their possibly high conformational flexibility [8], which motivates efforts to understand and predict their conformational landscape.

Structure prediction of polypeptides with amino acid length up to 40 amino acids in aqueous solution can be performed by a series of methods including deep-learning approaches such AlphaFold2 [9], TrRosetta [10], and APPTEST [11]. Looking at AlphaFold2, which revolutionized structure prediction of single folded domain to a root-mean-square deviation (RMSD) accuracy of 0.2 nm, its capability lies on machine learning based on protein data bank (PDB) [12] templates, multiple sequence alignments, co-evolution rules and sophisticated algorithms to predict local backbone and side conformations, and side chain - side chain contact probability within distances bins. AlphaFold2 builds the protein by energy minimization using a protein-specific energy potential.

TrRosetta is basically similar to AlphaFold2. It builds the protein structure based on direct energy minimizations with a restrained Rosetta. The restraints include inter-residue distance and orientation distributions, predicted by a deep neural network. Homologous templates are included in the network prediction to improve the accuracy further.

APPTEST also uses machine learning on the PDB [12] structures with a chain length between 5 and 40 amino acids. APPTEST derives C*α*-C*α* and C*β*-C*β* distance restraints, and backbone dihedral restraints that are input of simulated annealing and energy minimization.

Other methods which are accessible by WEB-servers or can be downloaded include Rosetta [13], I-TASSER [14], PepStrMod [15] and PEPFOLD [15, 16, 17]. Rosetta is a fragment-assembly approach based on Monte Carlo simulation, a library of predicted 9 and then 3 residues, and a coarse-grained model [13], followed by all-atom refinement. I-TASSER (Iterative Threading ASSEmbly Refinement) is a hierarchical approach that identifies structural templates from the protein data bank (PDB) by multiple threading approaches, with full-length atomic models constructed by iterative template-based fragment assembly simulations [14].

The PEPstrMOD server predicts the tertiary structure of small peptides with sequence length varying between 7 to 25 residues. The prediction strategy is based on the realization that *β*-turn is an important and consistent feature of small peptides in addition to regular structures. Thus, the method uses both the regular secondary structure information predicted from PSIPRED and *β*-turns information predicted from BetaTurns. The structure is further refined with energy minimization and molecular dynamic simulations.

PEP-FOLD2 is a fast accurate structure peptide approach based on the prediction of a profile of the structural alphabet of 4-amino length along the amino acid sequence, and a chain growth method based on the coarse-grained sOPEP2 model followed by Monte Carlo steps. It should be noted that PEP-FOLD2 is not free of learning as it uses an Support Vector Machine predictor relying on multiple sequence alignment. Of practical interest, during the time of this study, we could not access the APPTEST and PepStrMod servers. Also, TrRosetta cannot be applied to sequences with < 10 amino acids.

Overall, all these methods generate good models for well-structured peptides at pH 7 in aqueous solution because most structures deposited in the PDB from nuclear magnetic resonance (NMR) and X-ray diffraction models in the PDB were determined at neutral pH and the PDB contains close to 200,000 structures as of October 30th, 2022.

These methods face, however, two current limitations: correct conformational ensemble sampling of intrinsically disordered peptides or proteins (IDPs) which lack stable secondary and tertiary structures, and accurate conformational ensemble prediction of natural peptides as a function of pH and salt conditions. The first issue has motivated the development of new force fields, such as CHARMM36m-TIP3P modified [18] and AMBER99-DISP [19] and many others [20]. The current approach to address the impact of pH variations is to perform your own extensive molecular dynamics and replica exchange molecular dynamics simulations at your desired pH. Alternatively one can use pH-replica exchange molecular dynamics in proteins using a discrete protonation method [21] or all-atom and coarse-grained continuous constant pH molecular dynamics (CpHMD) method [22, 23, 24]. Accurate and fast peptide structure prediction at different pH and salt conditions is the objective of the present study.

The organization of this paper is as follows. In section 2, we present an extension of the coarse-grained, sOPEP2, force field to integrate Debye-Hückel charge interactions as a function of pH and salt concentrations. In section 3, we first present the results of structure predictions of six poly-charged amino acids as a function of pH and compare them to experimental CD data, and the predicted models obtained by TrRosetta and AlphaFold2. Next, we present the trRosetta, AlphaFold2 and PEP-FOLD with and without Debye-Hückel protocols and the analysis methods. These charged polypeptides are particularly interesting to assemble the sOPEP2 interactions and the Debye-Hückel charge interactions. This is followed by the prediction of 15 ordered peptides, which have NMR structures from pH 2 to pH 8. We finish this section on the prediction of four peptides for which low-resolution experimental data and topological description are available. Finally section 4 summarizes our findings.

## 3 Methods

### 3.1 the sOPEP2 force field

The sOPEP2 potential, to be used in a discrete space, originates from the OPEP potential [25, 26] which uses and explicit represention of the backbone (N, H, C*α*, N and H atoms) and one bead for each side chain, whose position to C*α* and Van der Walls depend on the side chain type. It is expressed as a sum of local, nonbonded and hydrogen-bond (H-bond) terms, with all parameters described [27].

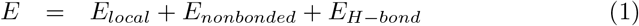

Since in PEP-FOLD the geometry is mainly imposed by the superimposition of the discrete SA letters, the local contributions are restricted to a simple flat-bottomed quadratic potential to described the energy associated with dihedral angles *ϕ* and *ψ*, described by:

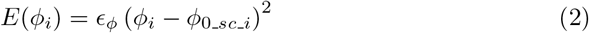

where *ϕ*_0_*sc_i*_ = *ϕ* within the interval [*ϕ_low_sc_i_, ϕ_high_sc_i_* and *ϕ*_0_*sc_i*_ = min(*ϕ* – *ϕ_low_sc_i_* – *ϕ_high_sc_i_*) outside of the interval *ϕ_low_sc_i_* and *ϕ_high_sc_i_* are specific to each amino acid type. (Binette et al. 2022)

Non bonded interactions corresponding to repulsion/attraction effects are described using the following Mie potential [28] given by:

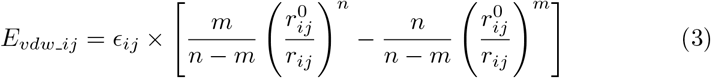

where *ϵ_ij_* is the potential depth and 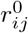 is the position of the potential minimum function of atomic types for *i* and *j*. The combination of exponents, *n* and *m,* gives the relationship between the position of the potential minimum (*r*^0^) and the position where it is zero (*gR*0):

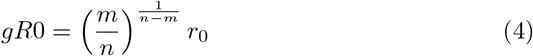

Hydrogen bonds are considered explicitly, using a combination of two types of contributions:

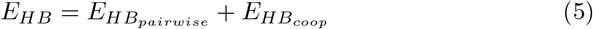

where *E_HB_pairwise__* corresponds to the pairwise contributions of hydrogen bond between residue (i) and residue (j), characterized by the hydrogen/acceptor distance *r_ij_* and the donor/hydrogen/acceptor angle *α_ij_*:

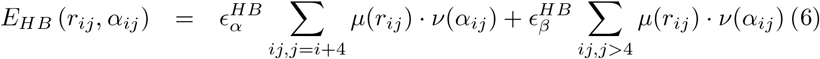

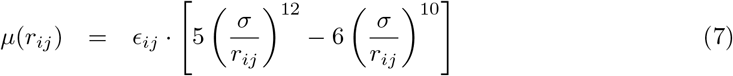

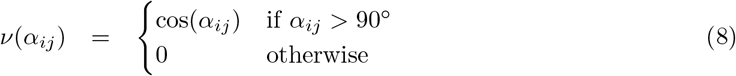

where *σ* = 0.18 nm is the position of the potential minimum and e is the potential depth. We distinguish between *α*-helix-like hydrogen bonds defined by O(i)-H(i+4) and other hydrogen bonds. Hydrogen bonds between a pair of residues separated by less than four amino acids are not considered.

*E_HB_coop__* involves pairs of hydrogen bonds (between residues (i) and (j) and residues (k) and (l)), so as to stabilize secondary structure motifs. The cooperativity energy is given by the following equations:

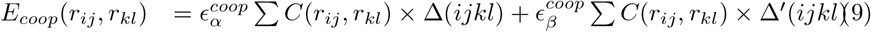

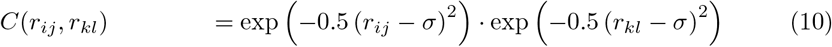

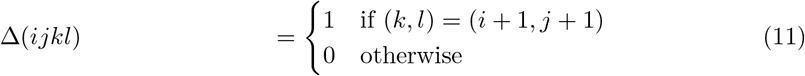

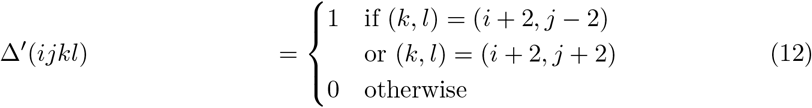

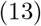

### 3.2 Debye-Hückel charge interactions

The new sOPEP version introduces the possibility to consider pH-dependent charge interactions, using the Debye-Hückel formulation [29].

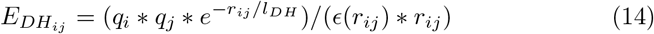

where *q_i_* and *q_j_* correspond to the charge of particles *i* and *j*, *j* > *i* + 1, respectively, *r_ij_* is the distance between the particles, *l_DH_* is the Debye length that depends of the ionic strength of the solvent, *ϵ*(*r_ij_*) is the dielectric constant that depends on the distance between the charges:

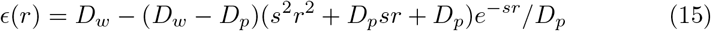

where *D_w_* is the dielectric constant of water, *D_p_* is the dielectric constant inside a protein, *s* is the slope of the sigmoidal function. In practice, we use values of 78, 2 and 0.6 for *D_w_, D_p_* and *s*, respectively, as stated in [30]

Since the sOPEP representation does not include all-atom side chains, but charges associated with particles of heterogeneous sizes, it is necessary to shift the energy curve to have energy values compatible with those of the Mie formulation. For each pair of particle, we shift the distance using:

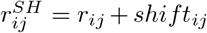 and we evaluate *E_DH_ij__* using 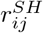 except for *ϵ*(*r*), where the unshifted distance is used.

Shift values are adjusted for r such as *E_vdw_r_* = *k, E_DH_r__* = *E_vdw_r_*, as illustrated Figure 1. In practice, we have found that values of k on the order of 4 kcal/mol are convenient. Finally, the Debye-Hückel energy is truncated to *E_DH_r__*, to avoid redundancy with the Mie potential.

**Figure 1:**
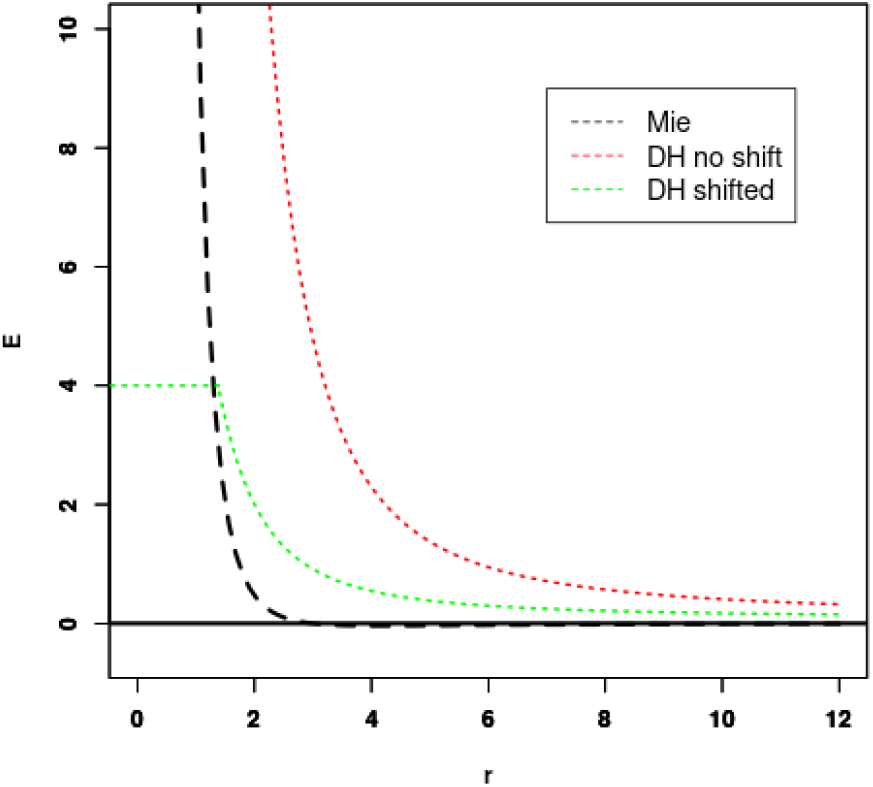
Fitting Debye-Hückel (DH) energy to Mie potential. The shift of the unshifted DH potential (red) is set so that the Mie (black) and shifted DH potential (green) cross for some energy threshold (4 kcal/mol in this case).

Finally, since sOPEP2 side-chain side-chain interactions already include some of the interaction between the charges, the Mie potential is restricted to only the repulsive part for charged side chains.

Charges are assigned to particles depending on the pH using pKa values of 3.9, 4.2, 6.0, 10.5 and 12.5 for ASP, GLU, HIS, LYS and ARG side chains, respectively, and 9. and 2. for N-terminal *α*-ammonium and C-terminal *α*-carboxyl groups, respectively. Note that it is possible to consider blocking the extremities using acetyl and N-methyl on N-terminus and C-terminus groups, respectively, in which case no charge is assigned to the extremity.

Finally, we have considered weighting differently the contributions depending on the separation of the amino acids in the sequence. In our experience, best results were obtained using a weight of 10 for residue separation of less than 7 and a weight of 2 otherwise.

### 3.3 PEP-FOLD, trRosetta and AlphaFold2 protocols and analysis

Our validation test set includes a total of 25 peptides as described in Section 4. For each peptide, we performed a total of 200 PEP-FOLD simulations, one TrRosetta simulation which uses PDB templates and homologous sequences, and one AlphaFold2 simulation in its standard version using 3 recycles, template information, and AMBER refinement. All methods, TrRosetta and AlphaFold2, PEP-FOLD without Debye-Hückel (referred to as PF-noDH), and PEP-FOLD with Debye-Hückel (PF-DH) return five models that we consider equiprobable. For PEP-FOLD results simulations, the fives models are based on their ranking using sOPEP2 total energy.

We have considered 15 peptides for which a PDB structure is available. These correspond to peptides previously studied during PEP-FOLD development and new peptides of structure released after September 1st, 2019, that are solved in pure aqueous environment. For the simulations of 15 peptides, the predicted models are evaluated by computing the CAD-score [31]. The reported CAD-score corresponds to the largest value of the cross CAD-scores between the five predicted models and all NMR structures. Following our previous work, if the CAD-score calculated on the backbone atoms is > 0.60, the model is associated with largely correct secondary structure prediction, otherwise if it is *>* 0.65 the model is correctly in terms of secondary and tertiary structures. For poly-charged amino acids, we also compute their secondary structure content using STRIDE program [32]. For the four sequences free of any NMR structure, we compare their predicted and experimental topologies.

## 4 Results and Discussion

secondary and tertiary structures. For poly-charged amino acids, we also compute their secondary structure content using STRIDE program [32]. For the four sequences free of any NMR structure, we compare their predicted and experimental topologies.

## 5 Results and Discussion

### 5.1 Predicted Models of poly-charged amino acids

For the simulations of the six poly-amino acids, namely (EK)15, (EK)5, (H)30, (E)15, (K)15 and (R)25, we calculated the alpha-helix, coil and turn contents averaged over all models and compare with circular dichroism (CD) experiments. It is to be noted that by default TrRosetta, AlphaFold2 and PF-noDH only perform simulations at neutral pH. CD values are not available at all pH varying from 3 to 13. We report, however, on the pH-dependent conformational ensemble using PEP-DH. Results are summarized in Table 1.

**Table 1:**
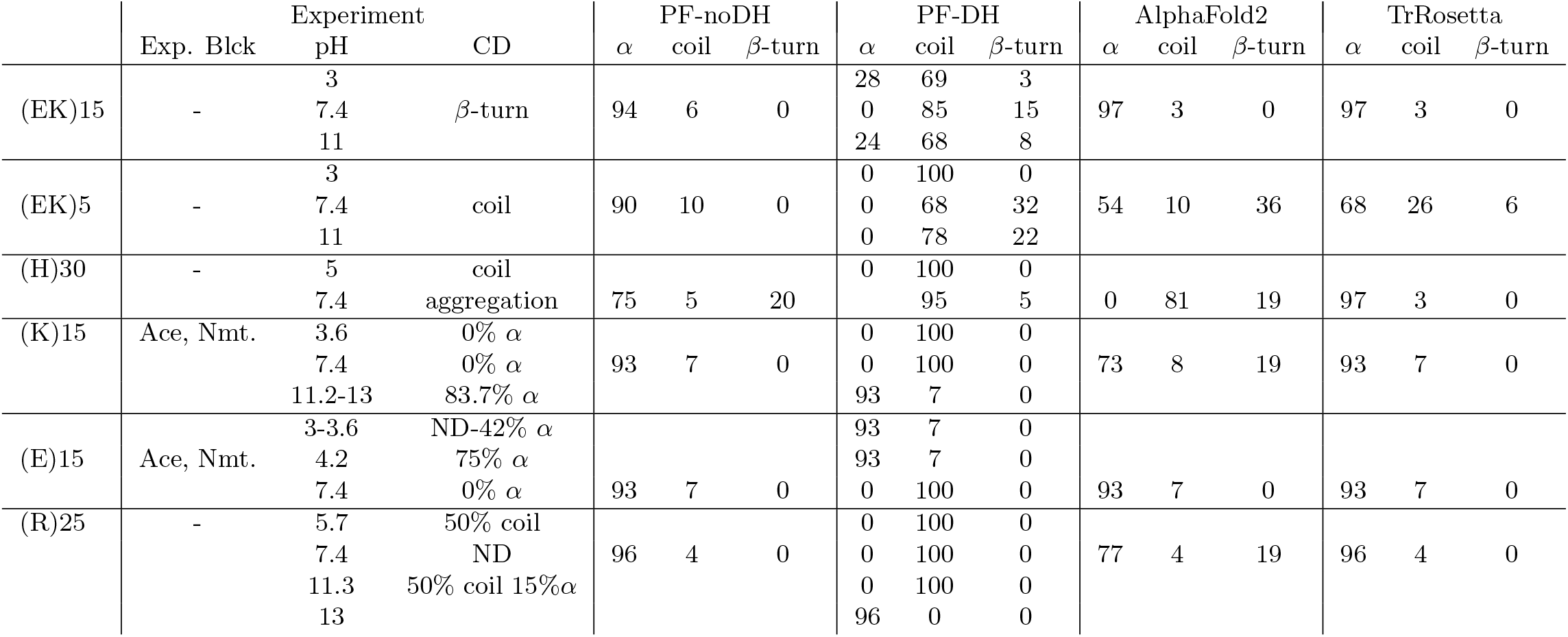
Structural impact of pH variation on polypeptides Results are presented for PEP-FOLD without and with Debye-Hueckel (PF-noDH, PF-DH), AlphaFold2 and trRosetta. Experimental information about the blocking of extremities (Exp. Bick.) using acetyl (Ace) or N-methyl (Nmt), pH and Circular Dichroism (CD) data, is reported. ND stands for not determined

TrRosetta, AlphaFold2, PF-noDH have a very high propensity to report alpha-helical conformations for the six polypeptides at pH 7.4, the exception being (H)30, with alpha-content varying from 54% to 97%, while CD displays only coil or beta-turn signals. For instance, for (EK)5, TrRosetta reports 68% helix and 26% coil, AlphaFold2 reports 54% helix and 10% coil and PF-noDH reports 90% helix and 10% coil. Only PF-DH is able to predict the CD coil character of (EK)5 with 68% coil and 32% turn.

PF-DH is the single method to predict 85% coil and 15% turn consistent at pH 7.4 with the beta-turn CD signal of (EK)15 [33], and PF-DH predicts 100% coil at pH 5 consistent with the coil CD signal of (H)30 [34]. There is strong indication that (H)30 polymerizes at pH 7.4 forming beta-sheets. At this pH, PF-noDH and TrRosetta predict strong helical conformations, while PF-DH and AlaphaFold2 predict a random coil, with contents of 95% and 81%, respectively.

The polypeptides (K)15 and (E)15 are particularly interesting because the alpha-helix content changes inversely with the pH. As observed by CD, the helical content of (K)15 increases with pH, while the helical content of (E)15 decreases with pH [35]. (K)15 have 0% helix at pH 3.6 and 83.7% coil at pH 11-13 by CD. PF-DH finds 0% helix at PH 3.6 and 93% coil at pH 11-13 (Figure 2). In contrast to (K)15, (E)15 (Figure 3) have 42% helix at pH 3.6 and 0% helix at pH 11-13 by CD. PF-DH finds 93% helix at pH 3.6 and 100% coil at pH 11-13.

**Figure 2:**
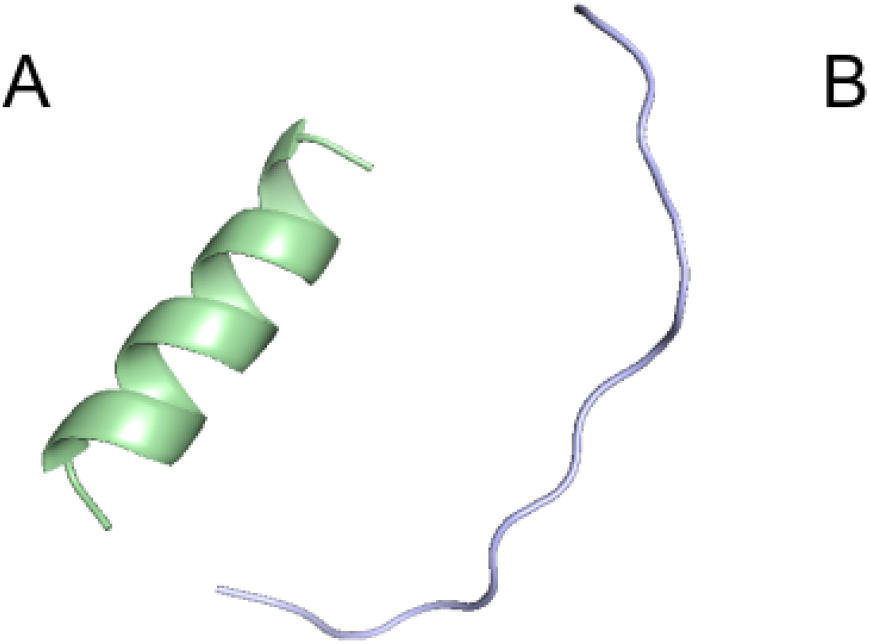
PF-DH Conformational ensemble of (K)15 as a function of pH. A: pH 7.4 B: pH 13. Only the lowest energy model is depicted.

**Figure 3:**
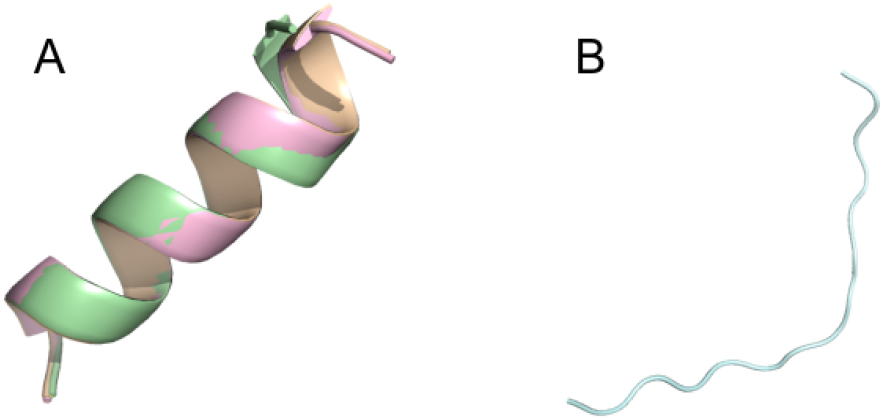
Conformational ensemble of (E)15 at pH7.4. A: PF-noDH, AlphaFold2, TrRosetta B: PF-DH. Only the lowest energy (rank 1) model is depicted.

The conformational ensemble of (R)25 is predicted to have 50% coil and 31% beta-sheet at pH 5.7 and have 51% coil and 21% beta-sheet at pH 11.3 by CD [36]. PF-DH predicts 100% coil, independently of the pH values. Its performance is however much better than those of PF-noDH, AlphaFold2 and TrRosetta which predict a high helical signal varying from 77% to 96%.

It is important to emphasize that in this study, we assume the standard pka values of charged amino acids irrespective of the amino acid composition of the peptides and the conformations of the peptides. This is a strong limitation of our current approach. Determining the pka values of charged amino acids in protein structures has motivated the development of many theoretical methods [21, 22, 23, 24]. To illustrate the variation of the pka values, we used the H++ server which is based on classical continuum electrostatics and basic statistical mechanisms [37]. Using (K)15, we found pka values ranging from 10.1 to 9.4 (versus 10.5 in our model); using (R)25, we found pka ranging from 9.6 to 11.6 in one conformation, and from 10.9 to 11.7 in another conformation (versus 12.5 in our model, a pH at which the peptide precipitates [36]), and using (H)30, we found pka variations from 4.7 to 6.3 (versus 6.0 in our model). Clearly this change of pka of the amino acids will impact the conformational ensemble of PF-DH.

### 5.2 Predicted Models of polypeptides with NMR structures

The experimental information of each well-ordered peptide, given in Table 2, include the amino acid length varying from 8 to 35 amino acids, the number of NMR models, the WDC (well-defined rigid core) according to the PDB, the topology, the pH, the ionic strength varying from 0 to 150 mM NaCl, the blocking of the extremities and the amino acid sequence.

**Table 2:**
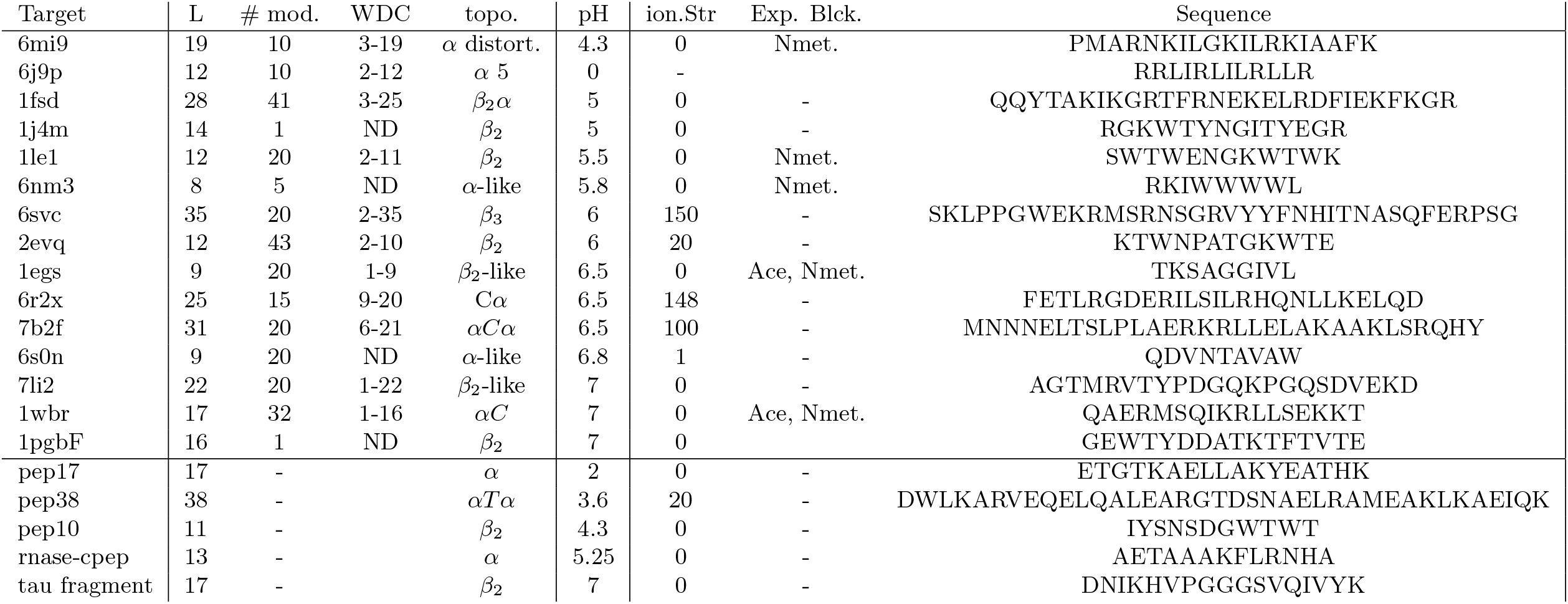
Peptide set For each peptide with an experimental structure available we specify its PDB identifier (PDB), size (L), the number of NMR models available (# mod.), its well defined core according to the PDB (WDC), its topology (topo.); and the experimental conditions including the pH, the ionic strenght (ion. Str.) and the presence of extra groups to block the N terminus (acetyl) and C terminus (N methyl) (Exp. Bick.), and the amino acid sequence. Four additional peptides without deposited structures but for which information exists in the literature are reported at the bottom of the table.

Table 3 reports on the CAD scores using the full sequences and the rigid sequence cores of each of the 15 peptides considered using the four methods.

**Table 3:**
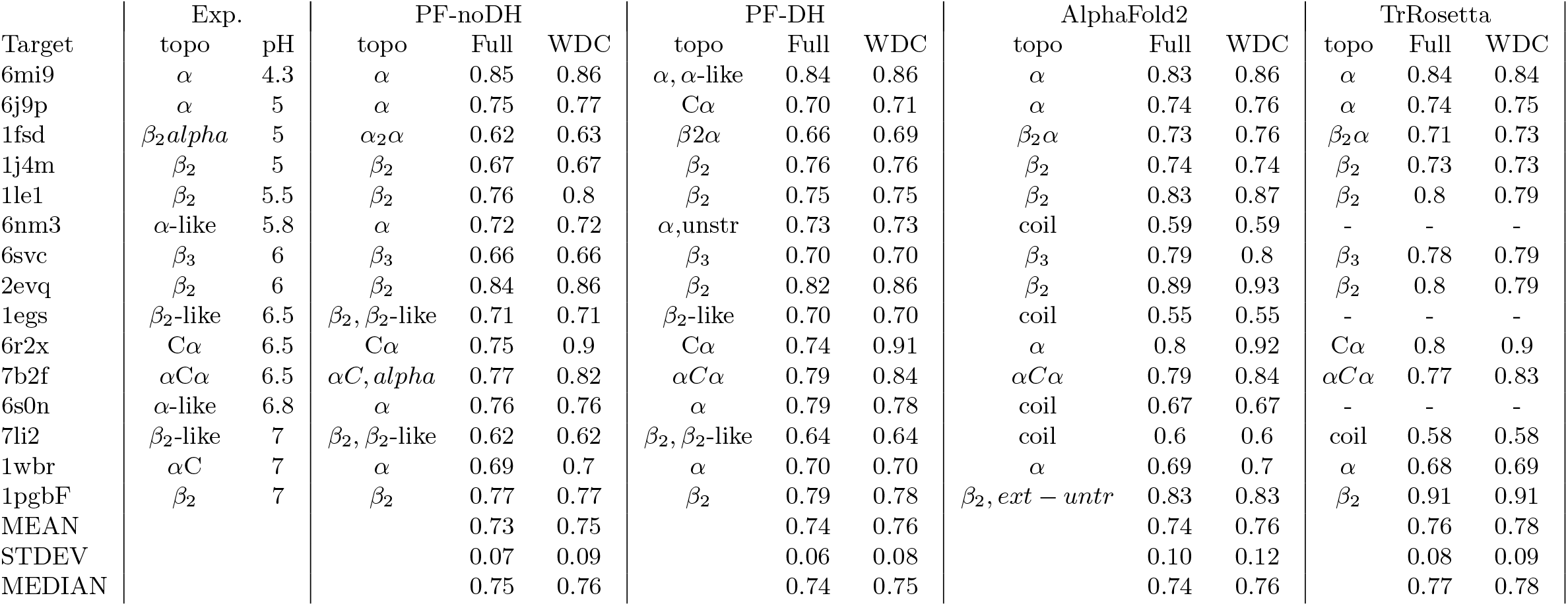
Performance prediction for structured peptides For each structure, we report a short description of the topology of the 5 best models, and the CADscore values (see methods) obtained for PEP-FOLD without and with Debye-Hückel (PF-noDH, PF-DH), AlphaFold2 and trRosetta. Note that TrRosetta is not functional for amino acid lengths < 10.

The first striking result is that (the mean, standard deviation and median) values of the CAD-scores averaged over the 16 peptides are nearly identical for the four methods using both the full sequences or the rigid cores. They reach (0.73, 0.07, 0.75) for PF-noDH, (0.74, 0.10, 0.74), for AlphaFold2, (0.74, 0.06, 0.74) for PF-DH and (0.76, 0.08, 0.77) for trRosetta using the full-sequences. Similar trends are observed considering the well defined core. Average CAD-scores being of 0.75, 0.76, 0.76 and 0.78 for PF-noDH, PF-DH, AlphaFold2 and trRosetta, respectively.

The second result is that PF-noDH and PF-DH do not predict any low quality models (CAD-score (< 0.6), while AlphaFold2 produces CAD-scores of 0.59, 0.55 and 0.6 for the three targets 6nm3,1egs and 7li2 (Figures 4, 5, 6, respectively), although these peptides are solved at neutral pH varying between 5.8 and 7. This low score results in differences between the experimental and predicted topologies. Experimentally, 6nm3 adopts a helical-like conformation, 1egs adopts a beta2-like conformation and 7li2 a beta-2 like conformation. For these three systems, AlphaFold2 predicts an extended-unstructured conformation. The 7li2 target is also problematic for TrRosetta, as it is the single system with a CAD-score <0.6, namely 0.58 leading to an extended-unstructured conformation. Inversely, TrRosetta is the best to predict the beta-hairpin of 1PGF [38] with a CAD-score of 0.91 versus 0.83 with AlphaFold2 and 0.79 with PF-DH.

**Figure 4:**
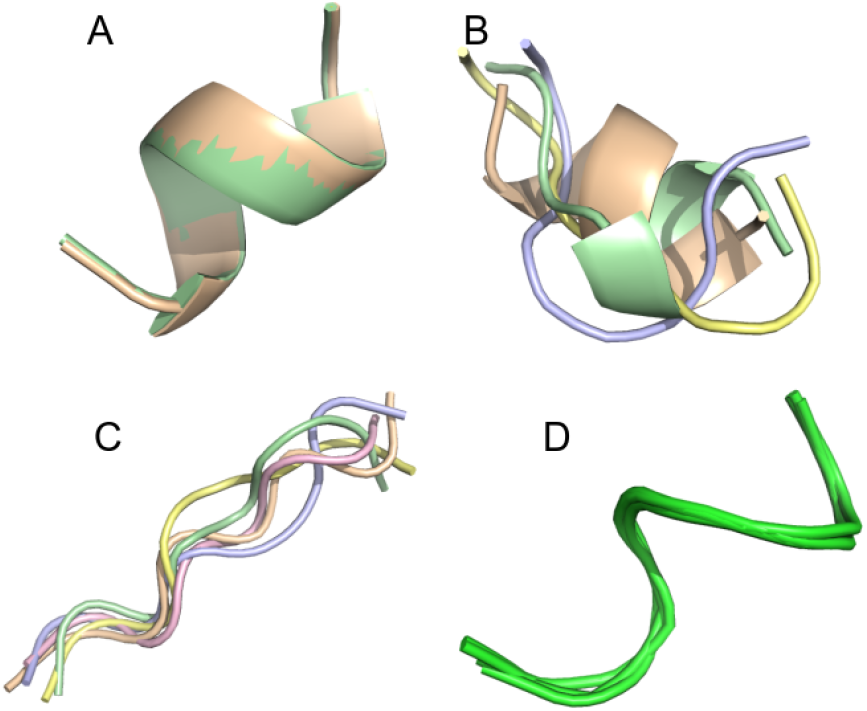
Conformational ensemble of 6nm3. A: PF-noDH, B: PF-DH at pH 4.3, C: AlphaFold2, D: NMR structure. For A, B and C, the 5 best models are depicted. For D, all models provided in the PDB are depicted.

**Figure 5:**
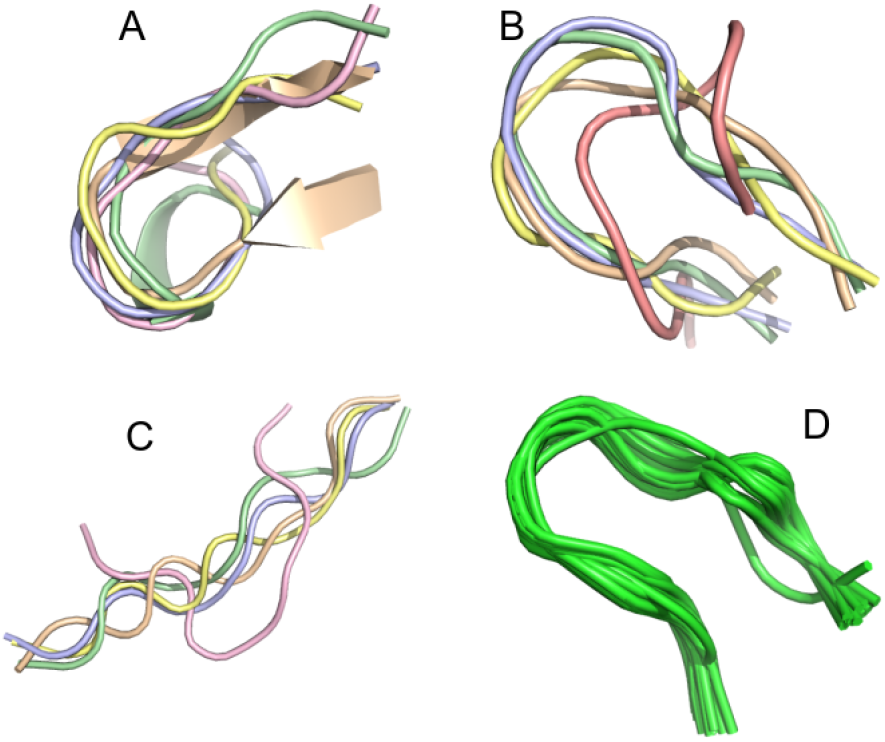
Conformational ensemble of 1egs. A: PF-noDH, B: PF-DH at pH 4.3, C: AlphaFold2, D: NMR structure. For A, B and C, the 5 best models are depicted. For D, all models provided in the PDB are depicted.

**Figure 6:**
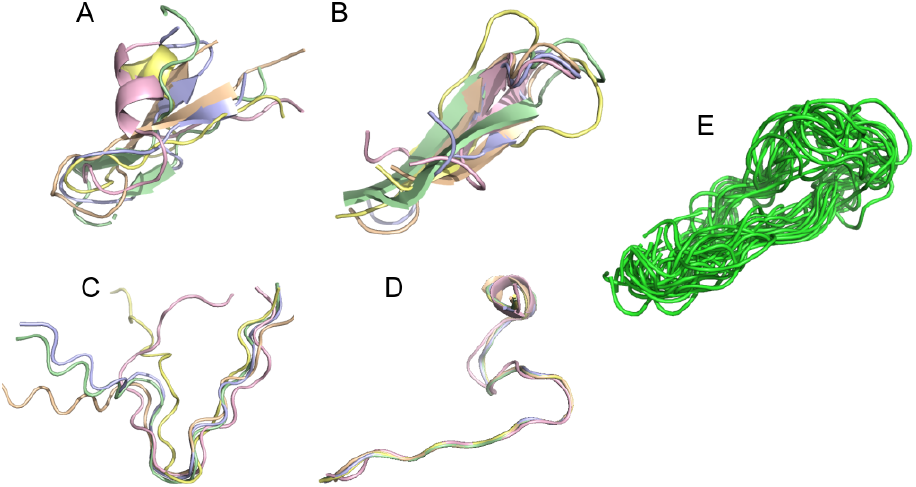
Conformational ensemble of 7li2. A: PF-noDH, B: PF-DH at pH 4.3, C: AlphaFold2, D:trRosetta, E: NMR structure. For A, B and C, the 5 best models are depicted. For D, all models provided in the PDB are depicted.

The third result is related to the performances of PF-DH with respect to PF-noDH, which provides evidence that the weights of the Debye-Hückel salt bridge interactions are consistent with the weights of sOPEP2 interactions. It was far from being evident that the addition of charged at extremities and charged amino acids in the core of the sequences would not change the quality of the models for pH varying between 2.9 to 7. The number of titratable amino acids vary from 1-2 (1le1, 1egs - 6nm3, 6evq), 4 for 1j4m, 5 for 6j9p, 6 for 6mi9, 7 for 1wbrand 7li2, 9 for 6r2x, 10 for 6svc and 7b2f, and 12 for 1fsd.

### 5.3 Predicted Models of polypeptides without any NMR structures

The last four peptides have been discussed in literature in terms of topological features without delivering any NMR structure. Their sequences are given at bottom of Table 2.

Two peptides are rather well described all four methods. Pep17 has been shown a stable monomeric helix at pH 2 using CD and NMR experiments [39]. PF-noDH, PH-DH at pH 2 and AlphaFold2 predict a helical conformatio with a frayed N-terminus, while TrRosetta predicts a full helical conformation (Figure 7). Pep38 determined experimentally by a helix-turn-helix [40] is also well reproduced by the four methods.

**Figure 7:**
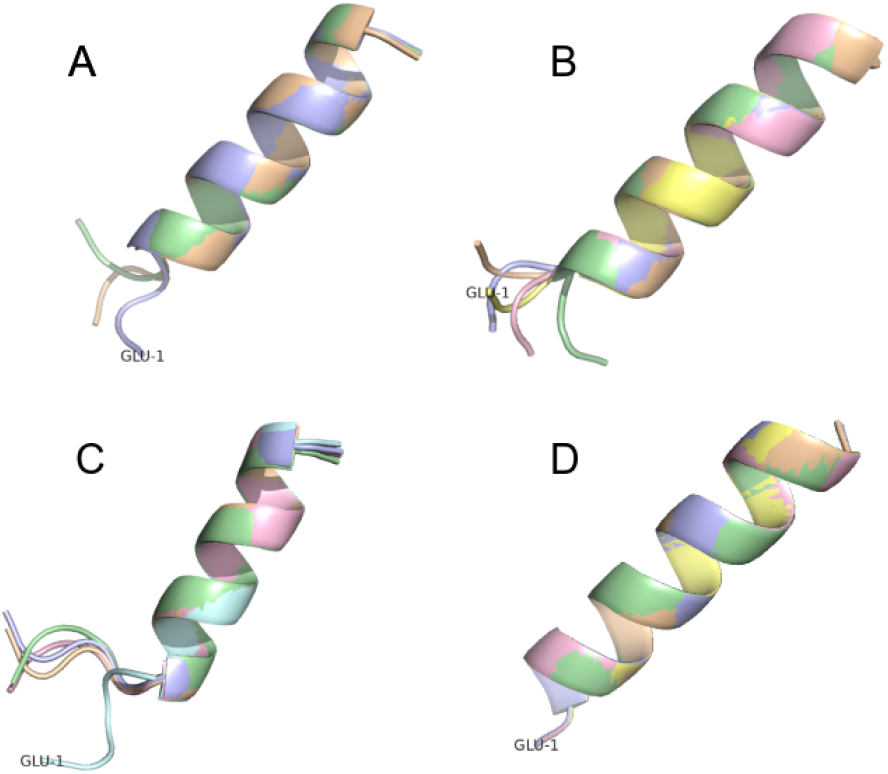
Conformational ensemble of pep17. A: PF-noDH, B: PF-DH - pH 2, C: AlphaFold2 and D: TrRosetta. For each, the 5 best models are depicted.

There are two cases, where AlphaFold2 and TrRosetta fail to produce the experimental data. The first peptide is pep10 which is described experimentally by an ensemble of distinct transient beta-hairpins [41]. It is described as an unstructured turn-like by trRosetta (8D), and an ensemble of extended and beta2-like conformations by AlphaFold2 (8C). In contrast, PEP-noDH and PEP-DH predict well a beta-hairpin conformations (8A and B)

The second peptide is the tau-fragment 295-306 containing the aggregation-prone PHF6 motif (306-K311). Using cross-linking mass-spectrometry, ab initio Rosetta [42], and CS-Rosetta which leveraged available chemical shifts [43] for the tau repeat spanning residues 243-365, the tau fragment 295-306 was predicted as a beta-hairpin [44]. PD-noDH and PF-DH predict the same conformation (Figure 9A and B). In contrast, Alpha-Fold2 predicts extended conformations (9C), and surprisingly trRosetta finds a random coil formation (9D).

**Figure 8:**
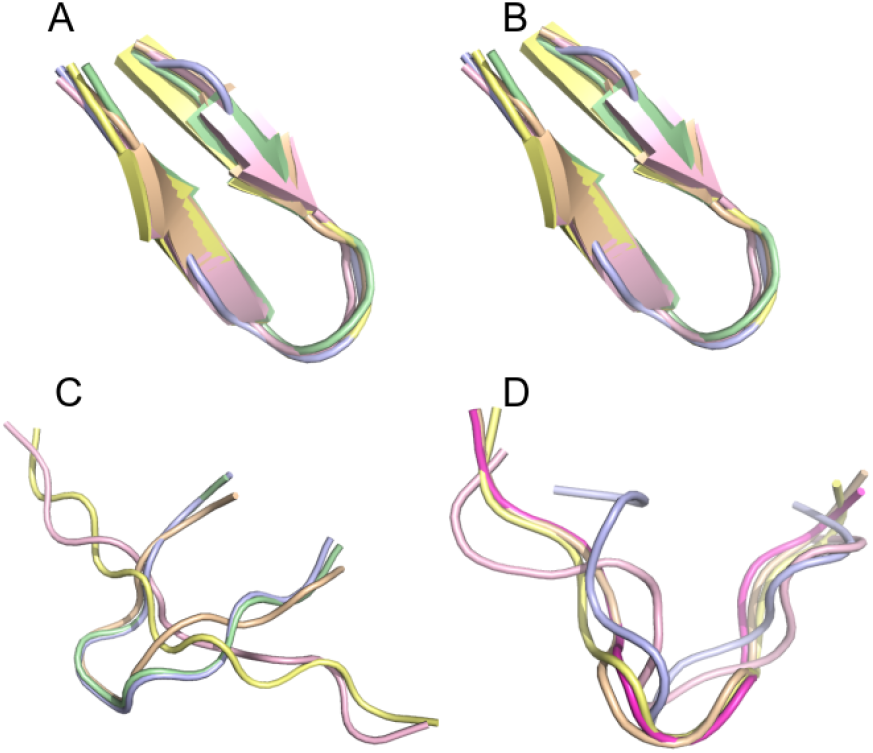
Conformational ensemble of pep10. A: PF-noDH, B: PF-DH at pH 4.3, C: AlphaFold2, D: TrRosetta. For each, the 5 best models are depicted.

**Figure 9:**
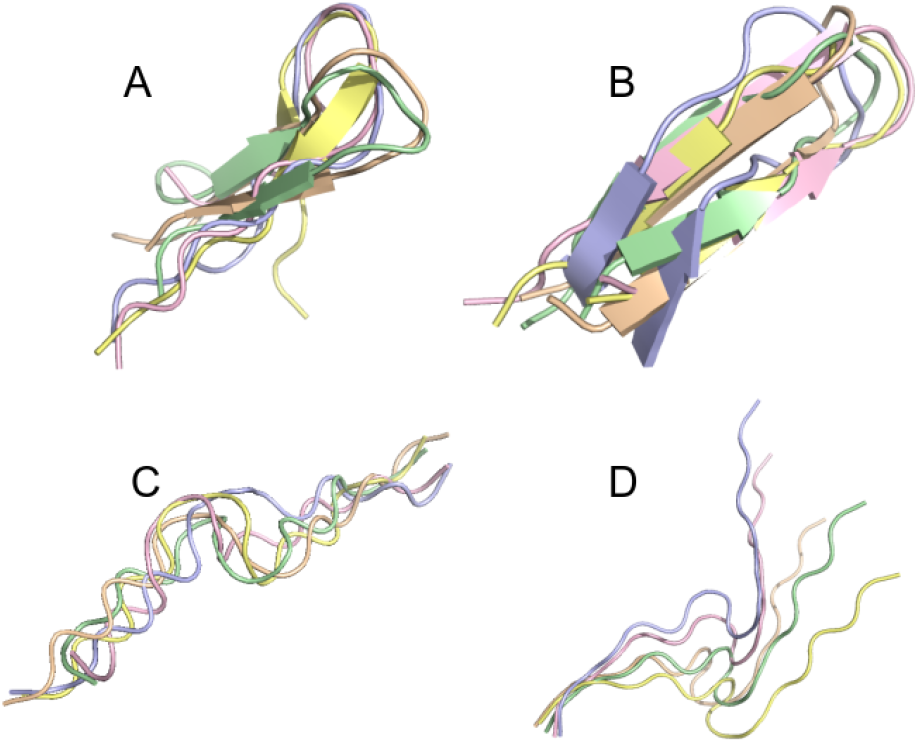
Conformation ensemble of tau-fragment at pH 7. A: PF-noDH, B: PF-DH, C: AlphaFold2 and D: TrRosetta. For each, the 5 best models are depicted.

Overall, this small set of peptides provides evidence of some limitations of AlphaFold2 and TrRosetta when the target does not have an homologous sequence in the PDB. By contrast, il also demonstrates for PEP-FOLD, that the learning stage of the local conformations that was performed from structures at neutral pH can be counterbalanced by the force field, making possible to explore new conformational regions depending on the pH. Finally, our new formulation also considers in principle the impact variations in salt concentration, but we could not identify from the literature any case reporting a conformational change upon ionic strength variation

### 5.4 Conclusions

Integrating pH variation effects to a coarse-grained model, where the side chains are represented by one single bead, is an important step toward accurate polypeptide structure prediction in aqueous solution, as coarse-graining with various granularities [45, 46], enhance sampling. This task has been performed by combining a Debye-Hückel formalism for charged - charged side chain interactions and the sOPEP2 potential. By using a total of 27 peptides of amino acid lengths varying between 7 to 38 amino acids, this study provides evidence that PF-FOLD, AlphaFold2 and TrRosetta performs similarly on peptides deposited in the Protein data Bank, but PF-FOLD outperforms the two recent machine-learning methods for poly-charged amino acids, and peptides for which homologous sequences are not deposited in the PDB.

Overall this work is one step towards peptide structure prediction in mimicking in vivo conditions. We are currently working on IDP’s in aqueous solution and de novo structure prediction of peptides at the surface of two-dimensional cell membranes.

## Acknowledgments

The authors thank A. Doig for helpful discussions about pKa.

## Notes

### Competing Interest Statement

The authors have declared no competing interest.

## References

[1] Sergueï O. Fetissov, Romain Legrand, and Nicolas Lucas. Bacterial Protein Mimetic of Peptide Hormone as a New Class of Protein-based Drugs. Current Medicinal Chemistry, 26(3):546–553, March 2019.

[2] Shira Ben-Shushan and Yifat Miller. Neuropeptides: Roles and Activities as Metal Chelators in Neurodegenerative Diseases. The Journal of Physical Chemistry B, 125(11):2796–2811, March 2021.

[3] Neeloffer Mookherjee, Marilyn A. Anderson, Henk P. Haagsman, and Donald J. Davidson. Antimicrobial host defence peptides: functions and clinical potential. Nature Reviews Drug Discovery, 19(5):311–332, May 2020.

[4] Milica Pavlicevic, Nelson Marmiroli, and Elena Maestri. Immunomodulatory peptides—A promising source for novel functional food production and drug discovery. Peptides, 148:170696, February 2022.

[5] Markus Muttenthaler, Glenn F. King, David J. Adams, and Paul F. Alewood. Trends in peptide drug discovery. Nature Reviews Drug Discovery, 20(4):309–325, April 2021.

[6] Walter Cabri, Paolo Cantelmi, Dario Corbisiero, Tommaso Fantoni, Lucia Ferrazzano, Giulia Martelli, Alexia Mattellone, and Alessandra Tolomelli. Therapeutic Peptides Targeting PPI in Clinical Development: Overview, Mechanism of Action and Perspectives. Frontiers in Molecular Biosciences, 8:697586, June 2021.

[7] Aviad Levin, Tuuli A. Hakala, Lee Schnaider, Gonçalo J. L. Bernardes, Ehud Gazit, and Tuomas P. J. Knowles. Biomimetic peptide self-assembly for functional materials. Nature Reviews Chemistry, 4(11):615–634, November 2020.

[8] Vasso Apostolopoulos, Joanna Bojarska, Tsun-Thai Chai, Sherif Elnagdy, Krzysztof Kaczmarek, John Matsoukas, Roger New, Keykavous Parang, Octavio Paredes Lopez, Hamideh Parhiz, Conrad O. Perera, Monica Pickholz, Milan Remko, Michele Saviano, Mariusz Skwarczynski, Yefeng Tang, Wojciech M. Wolf, Taku Yoshiya, Janusz Zabrocki, Piotr Zielenkiewicz, Maha AlKhazindar, Vanessa Barriga, Konstantinos Kelaidonis, Elham Mousavinezhad Sarasia, and Istvan Toth. A Global Review on Short Peptides: Frontiers and Perspectives. Molecules, 26(2):430, January 2021.

[9] John Jumper, Richard Evans, Alexander Pritzel, Tim Green, Michael Figurnov, Olaf Ronneberger, Kathryn Tunyasuvunakool, Russ Bates, Augustin Žídek, Anna Potapenko, and others. Highly accurate protein structure prediction with AlphaFold. Nature, 596(7873):583–589, 2021. Publisher: Nature Publishing Group.

[10] Zongyang Du, Hong Su, Wenkai Wang, Lisha Ye, Hong Wei, Zhenling Peng, Ivan Anishchenko, David Baker, and Jianyi Yang. The trRosetta server for fast and accurate protein structure prediction. Nature protocols, 16(12):5634–5651, 2021. Publisher: Nature Publishing Group.

[11] Patrick Brendan Timmons and Chandralal M Hewage. APPTEST is a novel protocol for the automatic prediction of peptide tertiary structures. Briefings in bioinformatics, 22(6):bbab308, 2021. Publisher: Oxford University Press.

[12] Peter W Rose, Chunxiao Bi, Wolfgang F Bluhm, Cole H Christie, Dimitris Dimitropoulos, Shuchismita Dutta, Rachel K Green, David S Goodsell, Andreas Prlić, Martha Quesada, and others. The RCSB Protein Data Bank: new resources for research and education. Nucleic acids research, 41(D1):D475–D482, 2012. Publisher: Oxford University Press.

[13] Richard Bonneau, Charlie EM Strauss, and David Baker. Improving the performance of Rosetta using multiple sequence alignment information and global measures of hydrophobic core formation. Proteins: Structure, Function, and Bioinformatics, 43(1):1–11, 2001. Publisher: Wiley Online Library.

[14] Yang Zhang. I-TASSER server for protein 3D structure prediction. BMC Bioinform., 9, 2008.

[15] Sandeep Singh, Harinder Singh, Abhishek Tuknait, Kumardeep Chaudhary, Balvinder Singh, S Kumaran, and Gajendra PS Raghava. PEP-strMOD: structure prediction of peptides containing natural, non-natural and modified residues. Biology direct, 10(1):1–19, 2015. Publisher: BioMed Central.

[16] Yimin Shen, Julien Maupetit, Philippe Derreumaux, and Pierre Tufféry. Improved PEP-FOLD approach for peptide and miniprotein structure prediction. Journal of chemical theory and computation, 10(10):4745–4758, 2014. Publisher: ACS Publications.

[17] Alexis Lamiable, Pierre Thévenet, Julien Rey, Marek Vavrusa, Philippe Derreumaux, and Pierre Tufféry. PEP-FOLD3: faster de novo structure prediction for linear peptides in solution and in complex. Nucleic Acids Research, 44(W1):W449–454, July 2016.

[18] Jing Huang, Sarah Rauscher, Grzegorz Nawrocki, Ting Ran, Michael Feig, Bert L De Groot, Helmut Grubmüller, and Alexander D MacKerell. CHARMM36m: an improved force field for folded and intrinsically disordered proteins. Nature methods, 14(1):71–73, 2017. Publisher: Nature Publishing Group.

[19] Paul Robustelli, Stefano Piana, and David E Shaw. Developing a molecular dynamics force field for both folded and disordered protein states. Proceedings of the National Academy of Sciences, 115(21):E4758–E4766, 2018. Publisher: National Acad Sciences.

[20] Phuong H. Nguyen and Philippe Derreumaux. Structures of the intrinsically disordered A*β*, tau and *α*-synuclein proteins in aqueous solution from computer simulations. Biophysical Chemistry, 264:106421, September 2020.

[21] Danial Sabri Dashti, Yilin Meng, and Adrian E. Roitberg. pH-Replica Exchange Molecular Dynamics in Proteins Using a Discrete Protonation Method. The Journal of Physical Chemistry B, 116(30):8805–8811, August 2012.

[22] Yandong Huang, Wei Chen, Jason A. Wallace, and Jana Shen. All-Atom Continuous Constant pH Molecular Dynamics With Particle Mesh Ewald and Titratable Water. Journal of Chemical Theory and Computation, 12(11):5411–5421, November 2016.

[23] Noora Aho, Pavel Buslaev, Anton Jansen, Paul Bauer, Gerrit Groenhof, and Berk Hess. Scalable Constant pH Molecular Dynamics in GROMACS. Journal of Chemical Theory and Computation, 18(10):6148–6160, October 2022.

[24] Fernando Luís Barroso da Silva, Samuela Pasquali, Philippe Derreumaux, and Luis Gustavo Dias. Electrostatics analysis of the mutational and ph effects of the n-terminal domain self-association of the major ampullate spidroin. Soft Matter, 12:5600–5612, 2016.

[25] Julien Maupetit, P. Tuffery, and Philippe Derreumaux. A coarse-grained protein force field for folding and structure prediction. Proteins: Structure, Function, and Bioinformatics, 69(2):394–408, November 2007.

[26] Fabio Sterpone, Simone Melchionna, Pierre Tuffery, Samuela Pasquali, Normand Mousseau, Tristan Cragnolini, Yassmine Chebaro, Jean-Francois St-Pierre, Maria Kalimeri, Alessandro Barducci, Yoann Laurin, Alex Tek, Marc Baaden, Phuong Hoang Nguyen, and Philippe Derreumaux. The OPEP protein model: from single molecules, amyloid formation, crowding and hydrodynamics to DNA/RNA systems. Chem. Soc. Rev., 43(13):4871–4893, 2014.

[27] Vincent Binette, Normand Mousseau, and Pierre Tuffery. A Generalized Attraction–Repulsion Potential and Revisited Fragment Library Improves PEP-FOLD Peptide Structure Prediction. Journal of Chemical Theory and Computation, 18(4):2720–2736, April 2022.

[28] Gustav Mie. Zur kinetischen Theorie der einatomigen Körper. Annalen der Physik, 316(8):657–697, 1903.

[29] Peter Debye and Erich Hückel. Zur Theorie der Elektrolyte. I. Gefrierpunk-tserniedrigung und verwandte Erscheinungen. Physikalische Zeitschrift, 24(185):305, 1923.

[30] Michio Iwaoka, Koji Yoshida, and Taku Shimosato. Application of a Distance-Dependent Sigmoidal Dielectric Constant to the REMC/SAAP3D Simulations of Chignolin, Trp-Cage, and the G10q Mutant. The Protein Journal, 39(5):402–410, October 2020.

[31] Kliment Olechnovič, Eleonora Kulberkytė, and Ceslovas Venclovas. CAD-score: A new contact area difference-based function for evaluation of protein structural models. Proteins: Structure, Function, and Bioinformatics, 81(1):149—162, January 2013.

[32] Dmitrij Frishman and Patrick Argos. Knowledge-based protein secondary structure assignment. Proteins: Structure, Function, and Genetics, 23(4):566–579, December 1995.

[33] Josh Smith, Patrick McMullen, Zhefan Yuan, Jim Pfaendtner, and Shaoyi Jiang. Elucidating Molecular Design Principles for Charge-Alternating Peptides. Biomacromolecules, 21(2):435–443, February 2020.

[34] Carlos Noble Jesus. On the Self-Assembly of pH-Sensitive Histidine-Based Copolypeptides. University College London, London, 2020.

[35] Piotr Batys, Maria Morga, Piotr Bonarek, and Maria Sammalkorpi. pH-Induced Changes in Polypeptide Conformation: Force-Field Comparison with Experimental Validation. The Journal of Physical Chemistry B, 124(14):2961–2972, April 2020.

[36] Maria Morga, Piotr Batys, Dominik Kosior, Piotr Bonarek, and Zbigniew Adamczyk. Poly-L-Arginine Molecule Properties in Simple Electrolytes: Molecular Dynamic Modeling and Experiments. International Journal of Environmental Research and Public Health, 19(6):3588, March 2022.

[37] R. Anandakrishnan, B. Aguilar, and A. V. Onufriev. H++ 3.0: automating pK prediction and the preparation of biomolecular structures for atomistic molecular modeling and simulations. Nucleic Acids Research, 40(W1):W537–W541, July 2012.

[38] Alessandro Barducci, Massimiliano Bonomi, and Philippe Derreumaux. Assessing the Quality of the OPEP Coarse-Grained Force Field. Journal of Chemical Theory and Computation, 7(6):1928–1934, June 2011.

[39] Erin K. Bradley, John F. Thomason, Fred E. Cohen, Phyllis Anne Kosen, and Irwin D. Kuntz. Studies of synthetic helical peptides using circular dichroism and nuclear magnetic resonance. Journal of Molecular Biology, 215(4):607–622, October 1990.

[40] Youcef Fezoui, Peter J. Connolly, and John J. Osterhout. Solution structure of *α*t*α*, a helical hairpin peptide of de novo design. Protein Science, 6(9):1869–1877, September 1997.

[41] Eva De Alba, Manuel Rico, and M. Angeles Jiménez. Cross-strand sidechain interactions versus turn conformation in *β*-hairpins. Protein Science, 6(12):2548–2560, December 1997.

[42] Sergey Ovchinnikov, Hahnbeom Park, David E. Kim, Frank DiMaio, and David Baker. Protein structure prediction using Rosetta in CASP12. Proteins: Structure, Function, and Bioinformatics, 86:113–121, March 2018.

[43] Oliver F. Lange, Paolo Rossi, Nikolaos G. Sgourakis, Yifan Song, Hsiau-Wei Lee, James M. Aramini, Asli Ertekin, Rong Xiao, Thomas B. Acton, Gaetano T. Montelione, and David Baker. Determination of solution structures of proteins up to 40 kDa using CS-Rosetta with sparse NMR data from deuterated samples. Proceedings of the National Academy of Sciences, 109(27):10873–10878, July 2012.

[44] Dailu Chen, Kenneth W. Drombosky, Zhiqiang Hou, Levent Sari, Omar M. Kashmer, Bryan D. Ryder, Valerie A. Perez, DaNae R. Woodard, Milo M. Lin, Marc I. Diamond, and Lukasz A. Joachimiak. Tau local structure shields an amyloid-forming motif and controls aggregation propensity. Nature Communications, 10(1):2493, December 2019.

[45] Sjoerd de Vries and Martin Zacharias. Flexible docking and refinement with a coarse-grained protein model using ATTRACT: Flexible Protein-Protein Docking and Refinement. Proteins: Structure, Function, and Bioinformatics, 81(12):2167–2174, December 2013.

[46] Adam K. Sieradzan, Cezary Czaplewski, Paweł Krupa, Magdalena A. Mozolewska, Agnieszka S. Karczyńiska, Agnieszka G. Lipska, Emilia A. Lubecka, Ewa Gołaś, Tomasz Wirecki, Mariusz Makowski, Stanisław Ołdziej, and Adam Liwo. Modeling the Structure, Dynamics, and Transformations of Proteins with the UNRES Force Field. In Victor Muñoz, editor, Protein Folding, volume 2376, pages 399–416. Springer US, New York, NY, 2022.

